# Where is God? A comparison of the neural correlates of mystical and religious praying

**DOI:** 10.64898/2026.02.22.707337

**Authors:** Sergio Elías Hernández, Katya Rubia, Oscar Perez-Diaz, José L. González-Mora, Alfonso Barrós-Loscertales

## Abstract

The perception of God can be as a transcendent entity that is infinite and outside of human beings, typical for religious traditions, or as an immanent entity that is outside and inside of human beings, typical for mystical traditions. These different perceptions of God may be associated with different neural correlates depending on which God we pray to. To elucidate the neural correlates of these different perceptions of the divine, we compared fMRI activation during praying between 18 Christians and 16 practitioners of Sahaja Yoga Meditation, characterised by transcendent and immanent perceptions of God, respectively. The thalamus was deactivated during praying in Meditators relative to Christians. Due to the sensory relay function of thalamus, the thalamic deactivation in meditators presumably reflects a reduction in the perception of external stimuli in order to focus on the internal perception of an immanent God, while the activation of the thalamus in Christian prayers could be associated with the dialogue with an externally perceived transcendent God.

## Introduction

The aim of prayers for people with religious or spiritual beliefs is to communicate with God. However, this communication is likely to be very different depending on how God is perceived. The underlying neuronal activity during prayers of different types of religious, spiritual and/or mystical practices may constitute a key factor to better understand the heterogeneity of the perception of the concept of God. The neuroscience research of prayers furthermore reveals valuable insights into how different spiritual practices activate our human brain [1–5].

Functional magnetic resonance imaging (fMRI) studies show that religious praying when God is perceived as a different and transcendent entity from oneself has similarities with a conversation with another person, activating brain regions involved in social cognition and theory of mind [5,6]. On the other hand, praying to God when God is perceived as immanent and part of one-self under the assumption that everything that exists is God or divine, as in mystical traditions, may well involve brain regions related to a different state of awareness, depending on how the praying persons perceives themselves in relation to God [7]

The perception of our ultimate reality within and without is very much modulated by our beliefs about ourselves and the world around us. Among believers, those who accept the existence of God, there are many different conceptions that are moulded by each particular religion, cultural heritage, mystical tradition or personal spiritual experiences, etc [8,9]

When it comes to perceiving God, there seems to be a dichotomy between religious and mystical traditions: God can be perceived as transcendent (religious) or as immanent (spiritual or mystical) [10]. The main attributes of a transcendent God, typical for western religions, typically are: God is separated and above His created universe. He has qualities beyond human capacities and human comprehension like omnipotence, omniscience and perfection, and there is a large difference between God who is Divine, infinite, perfect, unlimited and detached from the world and humans who are finite, imperfect, limited and not Divine. On the other hand, an immanent God, typical for mystical traditions such as Eastern Yoga, some Meditation practices, the Christian Gnosis and other mystical traditions, usually is characterized by His presence everywhere or omnipresence, as a part of human affairs and human history. The immanence of God is also associated with God’s nearness and accessibility to His/Her devotees. This immanence of God also implies that God is an essential part of our being and that human beings have a divine element to them. The devotee in his/her spiritual ascent finds the divine presence, the divine immanence in him/herself and all things by placing his/her attention inside rather than outside, and by means of achievement of the mystical union with God, the “Yoga” (i.e., union) between the individual “Atman” (the individual spiritual self or consciousness) and the all-pervading “Brahman” (the all-pervading divine cosmic consciousness) [10,11]. In mysticism, in particular during the mystical union between the individual and cosmic consciousness, the boundaries between the individual self/consciousness and the all-pervading divine consciousness become blurred or fused and the divine is perceived as immanent in every part of the Creation.

Sahaja Yoga Meditation (SYM) is a mystical meditation tradition that considers God as immanent. The aim of SYM is to achieve the state of yoga (yoga = union), a mystical state in which there is a connection between the Divine inside and outside the human beings. Through meditation there is the subjective perception of a “divine energy” within, called “Kundalini” energy (which means coiled energy in Sanskrit) and which is conceived of as being part of the all-pervading divine energy that innervates the Universe [12,13]. During the process of Yoga, the practitioner achieves a different, altered state of consciousness called the fourth state of consciousness or Turiya Awastha, i.e., the state of thoughtless awareness or mental silence [14–17].

Several studies have shown that depending on the type of religion and the type of prayer (improvised or not), different brain regions are activated during praying, including: the medial prefrontal cortex, the temporo-polar cortex, the temporo-parietal junction, the precuneus and the thalamus [5,8]. The medial prefrontal cortex (mPFC) in particular has been shown to be activated during improvised prayer [3]. This region is associated with self-reflection, social cognition, and theory of mind - the ability to attribute mental and emotional states to others with whom we interact with. The temporopolar and the temporo-parietal junction, in combination with the mPFC, are also part of the network of theory of mind [18]. The activation of areas that mediate theory of mind during spontaneous prayers suggests that some types of praying may be neurofunctionally similar to social interaction. The temporo-parietal regions have been related to the sensed presence of a “Sentient Being” related to the presence of God [19,20]. The activation of the precuneus has been shown during some prayers and has been linked to self-referential processing during praying.

Our previous fMRI study was focused on praying activity within practitioners of SYM only. The study showed predominantly deactivation of bilateral thalamus that was specific to praying relative to secular speech [3]. Because the thalamus has an essential role in the perception and integration of external sensory and somatosensory stimuli, arousal and consciousness [21], the deactivation of the thalamus during praying was interpreted as an increase of internalised attention and a decrease of external attention to exterior stimulation or sensations during prayers as part of the internalised attentional process and the reduction of the interference of senses, which are both necessary to reach the state of mental silence. However, given that in that study we did not include a religious control group, we could not assess how this activation during the mystical praying activity in SYM differs from praying in a Christian context or whether the thalamic deactivation would also be observed in Christian prayers.

The aim of this study was therefore to include a control group of Christian prayers to compare the brain activation patterns of praying within the mystical context of SYM with the brain activation patterns of praying within the Christian tradition. While both mystical (SYM) and religious (Christian) traditions pray in the form of a dialogue with God, the mystical tradition in SYM tries to reach the state of mental silence which is perceived as a union between the individual self (or individual spiritual energy) and the divine (or all-pervading divine energy); while in the Christian tradition, the goal is to have a dialogue with an external divine entity that is far beyond human conception and human limitations. The goal of the comparison of praying within SYM compared to that of a non-meditating Christian population is hence to highlight the neurofunctional differences between mystical and Christian praying experiences that presumably have differential neurofunctional correlates due to the differential experience and perception of the divine (i.e. the meditator’s mystical experience of union with the immanent divine versus the Christian experience of a dialogue with an external transcendent divine entity).

Based on our previous fMRI research on the activation patterns during prayer within SYM Meditators only [3] and based on fMRI studies that tested only Christian prayers [5], we hypothesized that these different conceptions and/or perceptions of God and differential experience of dialogue with the divine would be associated with neuro-phenomenologically different brain activation patterns. In particular, given our previous fMRI findings of reduced thalamic activation during the recitation of prayers within SYM, we hypothesised that this thalamic deactivation pattern would be specific to SYM compared to the recitation of prayers within the Christian tradition.

## Materials and Methods

### Participants

Sixteen expert practitioners of SYM (age range 24-63 years; 10 females) and an equivalent group of 18 non-meditating Christian control subjects (age range 23-62 years; 11 females), volunteered to participate in this study. As can be seen in Table 1, all meditators had more than 7 years of daily practice experience in SYM, while Christian controls reported that they had never experienced formal meditation practice throughout their lives. All volunteers reported (in a 5-point Likert scale) their frequency of praying the Lord’s Prayer and Spontaneous prayers. Additionally, meditators reported their average time dedicated to meditation per day and the frequency of the perception of the state of mental silence. After the scan session, all participants were asked to assess the quality of their praying experience inside the scanner. They furthermore completed a questionnaire on religious beliefs and practices, including belief in God, confidence in God’s reciprocity and frequency of praying (see Table 1). None of the participants reported any physical or mental illness, history of neurological disorders, addiction to alcohol, nicotine or other drugs.

**Table 1.** Demographic, meditative and religious data characteristics of the groups.

| Measures | SYM Meditators<br>Mean (SD)<br>[Range] | Christian Controls<br>Mean (SD)<br>[Range] | *t* (*df* =<br>1;32) | *p*-<br>value |
| --- | --- | --- | --- | --- |
| Age (years) | 48.2 (10.3)<br>[25-63] | 48.8 (10.8)<br>[23-62] | 0.15 | 0.88 |
| Education degree, | 2.4 (1.6)<br>[1-6] | 2.8 (1.4)<br>[1-5] | 0.76 | 0.46 |
| Height (cm) | 168.6 (7.7)<br>[159– 183] | 170.8 (8.1)<br>[154-187] | 0.83 | 0.41 |
| Weight (kg) | 73.8 (12.8)<br>[54-109] | 82.2 (24.1)<br>[57-110] | 1.75 | 0.09 |
| Lord's Prayer, weekly frequency | 4.5 (3.8)<br>[0-14] | 5.1 (2.5)<br>[0-7] | 0.48 | 0.64 |
| Spontaneous Prayer, weekly frequency | 5.7 (3.6)<br>[3-17] | 8.5 (4.8)<br>[0-14] | 1.94 | 0.06 |
| Do you believe in God? (1 to 5) | 5 (0)<br>[5-5] | 4.9 (0.5)<br>[3-5] | -0.94 | 0.35 |
| Do you believe that Santa Claus exists? (1 to 5) | 2.1 (1.1)<br>[1-4] | 1.6 (1.2)<br>[1-5] | -1.28 | 0.21 |
| Confidence in God's reciprocity? (1 to 5) | 4.8 (0.5)<br>[3-5] | 4.6 (0.7)<br>[3-5] | -0.93 | 0.36 |
| Years of SYM | 19.6 (8.1)<br>[7-30] | – | – | – |
| Duration of daily Meditation (min) | 47.1 (25.0)<br>[20-120] | – | – | – |
| Frequency of mental silence perception (1 to 5) | 3.6 (1.1)<br>[1-5] | – | – | – |

All participants signed a written informed consent form before study participation. The Human Research Ethics Committee of the University of La Laguna, Tenerife, Spain approved this study protocol (approval number: CEIBA 2011-0023), to protect the participants’ rights according to the rules of research at the University of La Laguna and according to the Declaration of Helsinki.

### Conditions and procedure

Following our previous fMRI paper that was based on Schjoedt’s fMRI study on prayer with Christian practitioners [3,5], a stimulus paradigm with five conditions was used. Four of the conditions followed a two-by-two factorial design between ‘Domain’ (spiritual vs. secular) and ‘Speech-act’ (formalised vs. improvised) factors (levels) (see Table 2). For the formalised and improvised spiritual speech-acts, the recitation of the Lord’s Prayer (henceforth, Lord’s_P) and Spontaneous prayer (henceforth, Spont_P) were used, respectively. For the secular speech acts, the recitation of a formalised well-known poem (henceforth, Poem) and making improvised wishes to Santa Claus (henceforth, Santa) were used. Similar to Schjoedt’s paradigm [5], a control condition in which participants were asked to count backwards from 100 was used rather than a resting state baseline condition to avoid task versus resting state related effects.

**Table 2.**
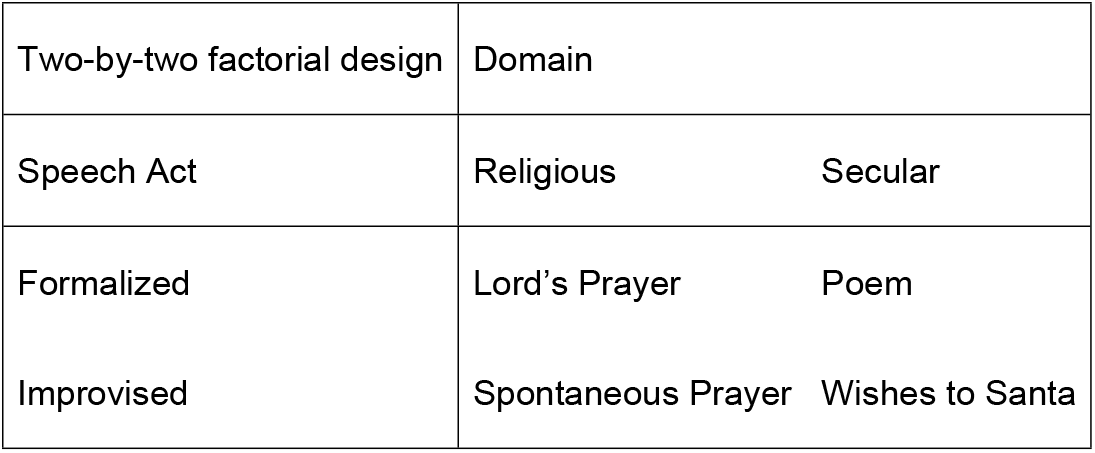
Factorial design two-by-two.

|  |  |  |
| --- | --- | --- |
| Two-by-two factorial design | Domain |  |
| Speech Act | Religious | Secular |
| Formalized | Lord's Prayer | Poem |
| Improvised | Spontaneous Prayer | Wishes to Santa |

First, participants underwent a 7 min structural scan where they were instructed to relax and prepare themselves for the posterior tasks. Afterwards, during the fMRI scan, volunteers were asked to perform the above mentioned five tasks in a pseudo-randomized order. Each task was preceded by their respective two seconds auditory instruction. Each of the five task conditions was repeated six times, each of them lasted 26s, excluding the 2s audio instruction. The total time of fMRI acquisition was 14min. During this time, tasks were performed silently as internal speech with eyes closed, and participants were asked to concentrate on the task at hand. They were instructed to repeat the current task if they finished it before the end of the 26s block.

### MRI Acquisition

Axially oriented BOLD functional images were obtained by a 3T Sigma HD MR scanner (GE Healthcare, Waukesha, WI, USA) using an echo-planar-imaging gradient-echo sequence and an 8-channel head coil with the following parameters: TR = 2000 ms, TE = 21,6 ms, flip angle = 90°, matrix size = 64 × 64 pixels, 37 slices, 4 × 4 mm in plane resolution, slice thickness = 4 mm, interleaved acquisition. The head was stabilized with foam pads. The slices were aligned to the anterior commissure—posterior commissure line and covered the whole brain. Functional scanning was preceded by 20 s of dummy scans to ensure tissue steady-state magnetization. A total of 420 fMRI whole brain volumes were taken during each participant’s single run.

For the structural acquisition before the fMRI acquisition, high-resolution sagittal oriented anatomical images were collected for anatomical reference; for this purpose, a 3D fast spoiled-gradient-recalled pulse sequence was obtained with the following parameters: TR = 8.844 ms, TE = 1.752 ms, flip angle = 10°, matrix size = 256 × 256 pixels, slice thickness = 1 mm.

### Data preprocessing and analysis

The functional brain images were pre-processed and analysed using Statistical Parametric Mapping, SPM12 (Wellcome Trust Centre for Neuroimaging, London United Kingdom). A general linear model was implemented for each participant at the first level effects, and the results were computed as a group-level effect (second-level effects). Pre-processing steps included: 1) within-subject registration and unwarping of time series, the realignment part of this routine realigns a time-series of images acquired from the same subject using a least squares approach and a 6 parameter (rigid body) spatial transformation; 2) co-registering individual structural images to the mean functional image of each subject; 3) co-registered images were segmented into grey matter, white matter, cerebrospinal fluid, bone, soft tissue and air/background; 4) spatial normalization of functional volumes by using the parameters extracted from the anatomical segmentation procedure in each subject and resampling voxel size to 2×2×2 mm; and 5) spatial smoothing with an 8-mm full-width-at-half-maximum (FWHM) Gaussian kernel, a high pass filter was used to remove low-frequency drifts in the fMRI BOLD signal. Only one subject presented extensive head movement (more than 3 mm/3º), but movement was appropriately corrected. We tested whether the exclusion of this participant affected the group statistics in a subgroup analysis. This was not the case and the subject was hence included in the analyses.

The first level analysis included 6 conditions: baseline condition (counting backwards), the audio instruction of each task (which was not used for posterior analysis), and the four conditions of interest: Santa, Poem, Lord’s_P and Spont_P.

The contrasts of interest computed were as follows: (1) Domain main effect (Spont_P & Lord’s_P > Santa & Poem and vice versa); (2) Speech act main effect (Poem & Lord’s_P > Santa & Spont_P and vice versa). Next, we tested for interactions (Spont_P & Poem > Lord’s_P & Santa and vice versa), simple effects (comparing Spont_P, Lord’s_P, Santa and Poem with each other), and simple contrasts comparing each condition with the base-line condition (Count). Baseline conditions was modelled implicitly. Additionally, this analysis included the head motion parameters as regressors of no interest in the model. Group analysis was performed by using the random-effect approach with a two-sample t-test. Analyses were thresholded at a statistical voxel-wise p <.001 and at corrected cluster p-value with multiple correction p <.05 FWE.

## Results

After the scanner sessions, all volunteers reported that they could properly concentrate and follow the tasks inside the scanner. In the post-scanner survey volunteers were asked to evaluate their performance for the different tasks from 5 “very well” to 1 “very bad”; on average, volunteers evaluated their performance between “well” (4) and “very well” (5), see detailed performance scores in Table 3.

**Table 3.** Post-Scanner Volunteer’s self-evaluation scores of their performance for each task.

| Task | SYM Meditators Mean (SD) | Christian Controls Mean (SD) |
| --- | --- | --- |
| Lord's Prayer | 4.5 (0.73) | 4.84 (0.37) |
| Spontaneous Prayer | 4.63 (0.62) | 4.74 (0.93) |
| Wishes to Santa | 4.13 (0.72) | 4.47 (0.90) |
| Poem | 4.38 (0.62) | 4.95 (0.23) |
| Counting backwards | 4.28 (0.82) | 4.79 (0.42) |

### Main effects of group differences in prayer-related activation

Because the results of the different prayer conditions in practitioners of SYM are already published [3], our primary objective was to examine group differences between meditators of SYM and non-meditating Christian practitioners.

The comparison between groups of the effects on the two prayer conditions relative to their secular contrasts: Spont_P & Lord’s_P > Santa & Poem showed a decrease in the neuronal activation of the bilateral thalamus in SYM compared to Christian controls (see Figure 1, Table 4).

**Table 4.**
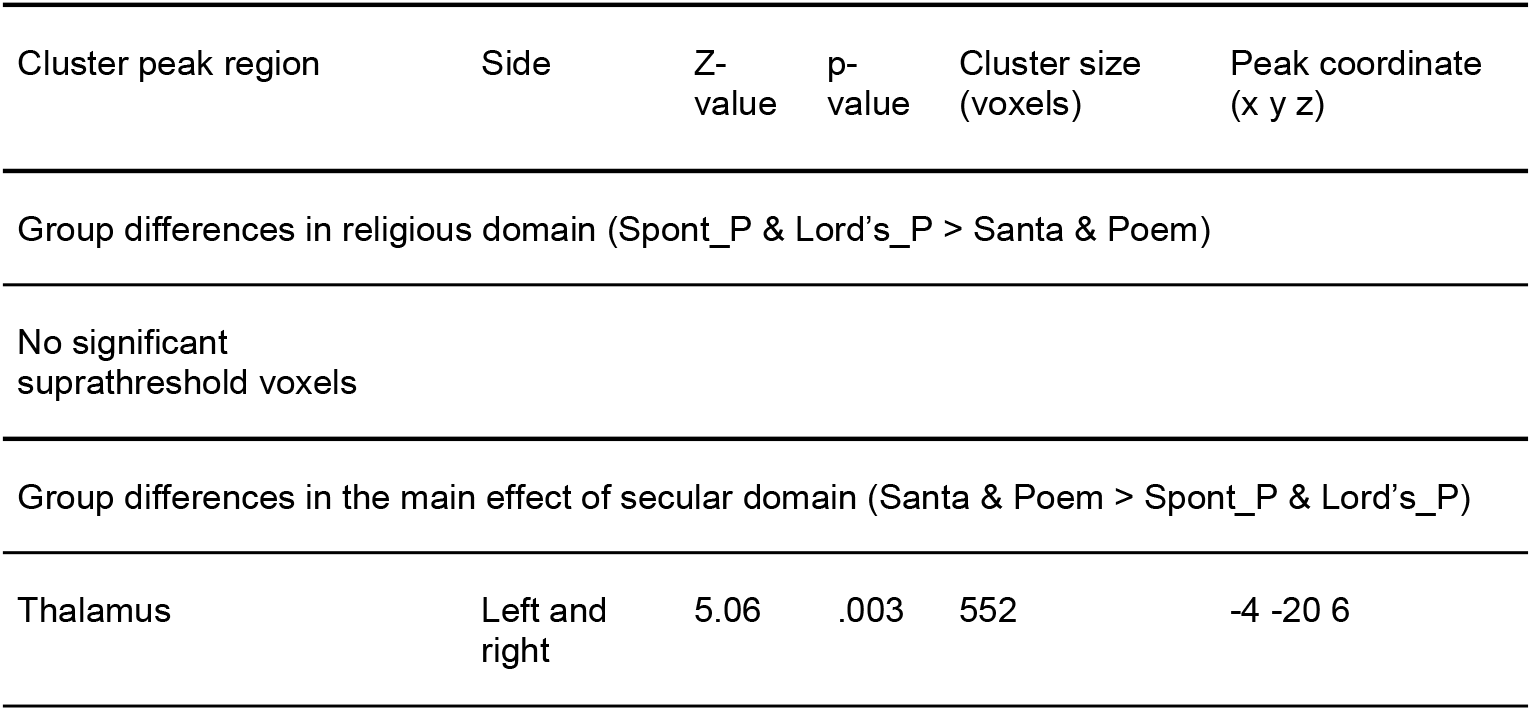
Two Sample t-Test between Christian controls > SYM Meditators during prayer-related activation

| Cluster peak region | Side | Z-value | p-value | Cluster size<br>(voxels) | Peak coordinate<br>(x y z) |
| --- | --- | --- | --- | --- | --- |
| Group differences in religious domain (Spont_P & Lord's_P > Santa & Poem) |  |  |  |  |  |
| No significant<br>suprathreshold voxels |  |  |  |  |  |
| Group differences in the main effect of secular domain (Santa & Poem > Spont_P & Lord's_P) |  |  |  |  |  |
| Thalamus | Left and<br>right | 5.06 | .003 | 552 | -4 -20 6 |

**Fig. 1.**
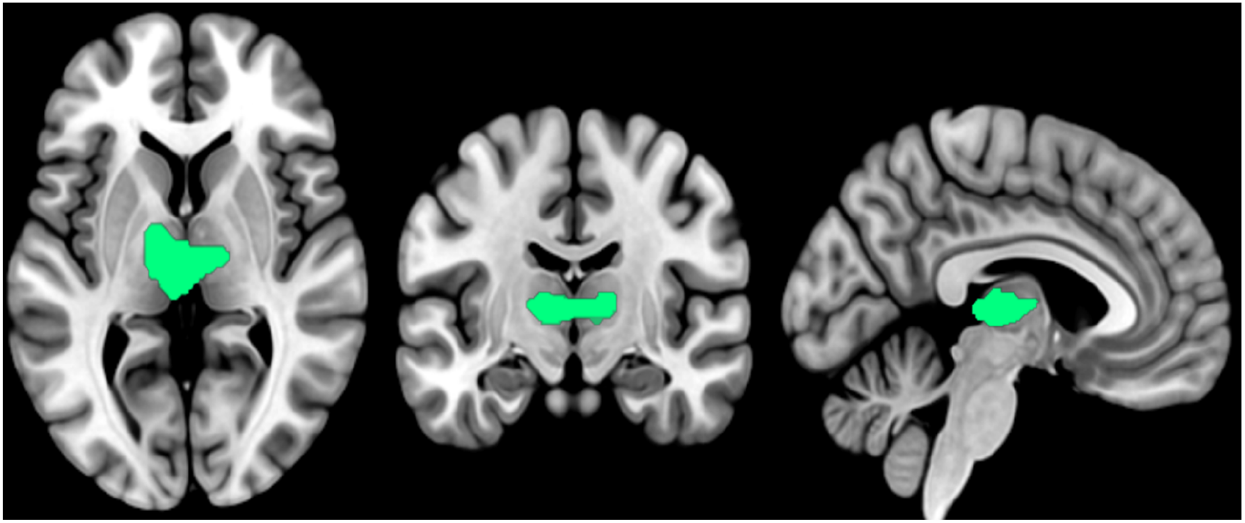
Group differences in the main effects of secular versus religious domain (Santa & Poem > Spont_P & Lord’s_P). Activation is decreased in the left and right thalamus during SYM versus Christian praying.

All the other comparisons between groups did not show any significant differences. Specifically, those comparisons that did not show significant differences between groups were: “Speech act” formalised versus improvised (Lord’s_P & Poem > Spont_P & Santa), the interactions between the two factors “domain” religious or secular (Lord’s_P & Santa > Spont_P & Poem), “Speech act” formalised versus improvised (Lord’s_P & Santa > Spont_P & Poem). No significant cluster survived any of the simple contrasts like: Spont_P > Santa, Lord’s_P > Poem or Lord’s_P > Spont_P.

## Discussion

The study showed that SYM practitioners compared to Christians showed reduced activation in bilateral thalamus during praying. The findings extend our previous finding that the thalamus is deactivated during prayers in SYM compared to non-praying by showing that the deactivation is specific to praying in SYM relative to Christian praying.

The thalamus mediates sensory and somatosensory processing of external stimuli and has an essential role in arousal [22], which is likely reduced in SYM when practitioners engage in prayer-related activity. The use of Christians as a comparison group may have enhanced the contrast between an internalized vs. an externalized divine agency. The deactivation of the thalamus in SYM practitioners during praying could be interpreted as a neural marker of an immanent focus on a mystical divine dialogue during meditation. It thus seems that the thalamus as a sensory relay station, receiving information from the senses (except smell) and directing it to the appropriate areas of the cerebral cortex for further processing [22], may be reduced in its activity in SYM favouring inner concentration by reduced sensory gating of external stimuli in the search of the immanent experience of God as an inner communication with the part of God that is internal.

Our previous fMRI study of functional connectivity related to SYM showed that during SYM, the thalamus was reduced in its functional connectivity to medial prefrontal cortex and adjacent anterior cingulate cortex and that this was associated with the state of mental silence [17]. This could suggest that in SYM during their mystical practices, the mPFC/ACC modulates this reduction of activation in the thalamus in the search of mental silence, aligning with the notion of an immanent divine conception (e.g. “God within”). This inner attention in SYM seems to be also reflected in the activation of the right anterior insula during SYM meditation [16] which is a key region of interoception and inner consciousness [23–25].

The thalamus controls perception of the external world [26–30] and modulates arousal, attention, cognition and consciousness [31–33], which is altered during meditation [34–36]. Recent work emphasizes the thalamus within its thalamocortical circuitries as a controller of large-scale network topology, enabling shifts between internally and externally oriented brain states that underpin consciousness [37]. Resting state fMRI studies showed that weaker functional connectivity between the thalamus and posterior cingulate, a core part of the default mode network reflecting mind-wandering, is associated with high traits of mindfulness [38] and with meditation-induced reduction of mind-wandering in patients with depression [39]. These findings suggest that the thalamus acts as a switch between mind-wandering and meditation, the latter of which has been shown to reduce mind-wandering [39]. The findings of thalamic deactivation are also analogous to findings of reduced functional connectivity between thalamus and primary sensory areas during Taoist Meditation [40] which may reflect greater inward focus and withdrawal of attention from the external world, associated with meditation [15]. Functional connectivity studies thus show that Meditation can modulate thalamus-default mode network and thalamus-sensory prefrontal coupling, which may underlie the experience of altered states of consciousness such as mental silence as experienced in SYM.

The thalamus is crucially implicated in both self-awareness and the awareness of the sensory environment (i.e., self-consciousness) via connections to brainstem arousal systems by forming a binding site to facilitate multisensory integration with the internal representations of self [41,42]. The reduction in thalamic activation may hence reflect volitionally mediated disengagement from self-referential appraisals of ascending sensory and nociceptive inputs to promote a sensory-self-centric decoupling mechanism [43]. This process would be in line with the concept of Meditation fostering a non-reactive sense of self to alleviate suffering [43]. This would be also in line with evidence for pain reduction with Meditation via insular and anterior cingulate modulation [44] as well as via reduced thalamic and cortical responses to nociceptive input [34].

In conclusion, meditative mystical versus religious praying reduces thalamic activation, presumably reflecting reduced coupling to cortico-subcortical external sensory and internal mind-wandering processes in order to focus on internal mystical experiences of inner consciousness leading to the state of mental silence.

## Acknowledgements

We acknowledge the support of MRI services for Biomedical Studies (Servicio de Resonancia Magnetica para Investigaciones Biomedicas) of the University of La Laguna. KR is supported by grants from the National Institute of Health Research (NIHR) (NIHR130077 and NIHR203684), by the NIHR Biomedical Research Centre at South London and Maudsley NHS foundation Trust and King’s College London (NIHR BRC Maudsley), the Medical Research Council (MRC) (APP32868), Medical Research Foundation (MRF-176-0002-RG-FLOH-C0929) and Rosetrees Foundation (3442198). The views expressed in this publication are those of the authors and not necessarily those of the National Health Service (NHS), the NIHR or the UK Department of Health and Social Care.

We warmly thank all the volunteers for their participation in this study.

